# Collection and preservation of urinary proteins using a fluff pulp diaper

**DOI:** 10.1101/130955

**Authors:** Weiwei Qin, Zhenhuan Du, Youhe Gao

## Abstract

Change is the most fundamental property of biomarker. Contrast to the blood, which is under homeostatic controls, urine reflects changes in the body earlier and more sensitive therefore is a better biomarker source. And drawing blood from infants and toddlers is hard and less tolerated. For patients limited by language, giving chief complaint is difficult. Thus, monitoring biomarkers in urine can provide valuable clues for diagnosis of diseases, especially pediatric diseases. Collecting urine from young children and some adult patients is more challenging than collecting it from healthy adults. Here, we propose a method that uses a fluff pulp diaper to collect urine. Urinary proteins were then eluted and adsorbed onto a piece of nitrocellulose membrane, which can be dried and stored in a vacuumed bag. SDS-PAGE and LC-MS/MS analysis indicated that this method is reproducible, and similar proteins were identified as those obtained using an acetone precipitation method. With this simple economical method, it is possible to collect and preserve urine samples from infants, toddlers, and patients with special needs, even for large-scale biomarker studies.

## Introduction

Biomarker is the measurable change associated with a physiological or pathophysiological process (Gao, 2013), its nature is change. In stark contrast to the blood, which is controlled by homeostatic mechanisms, urine as the waste of body accumulates changes (Gao, 2013; Li et al., 2014). Urine is a sensitive matrix affected by many factors, such as physiological conditions, age, gender, hormones, and diseases (Wu and Gao, 2015). Even brain diseases can be reflected in the urine (An and Gao, 2015). In addition, urine can easily be collected non-invasively in large quantities, and compared to other biofluids it does not undergo significant proteolytic degradation (Stéphane et al., 2008). We propose that urine is a better resource for biomarker research than blood.

Hundreds of candidate biomarkers associated with a wide range of diseases have been reported in urine (Shao et al., 2011). Thus, as a valuable biofluid, urine should be preserved comprehensively along with the patient’s medical record. This is a critical step for validation, which facilitates biomarker research and its translation from the bench to the bedside. We developed urimem that adsorb biological molecules in urine onto a membrane (Jia et al., 2014; Zhang et al., 2015), which makes preservation of urine samples economically feasible for large-scale and long-term research.

For patients limited by language ability, giving chief complaint is difficult. Diagnoses typically rely on laboratory examinations. However, blood sampling is invasive and has additional challenges with infants and toddlers. Monitoring biomarkers in urine can provide valuable clues for diagnosis of diseases, especially for children. However, collecting urine from young children and some adult patients is more complicated than collecting from healthy adults. Here, we propose a method using a fluff pulp diaper to collect and preserve urine.

## Results and Discussion

### SDS-PAGE analysis of urinary proteins

Urinary protein samples were separated using 12% SDS-PAGE. Samples prepared following the diaper method we have described reveal no protein degradation and isolation of the same bands as those from samples prepared by acetone precipitation, indicating that the diaper method has good technical reproducibility (Fig. 1).

**Figure 1.**
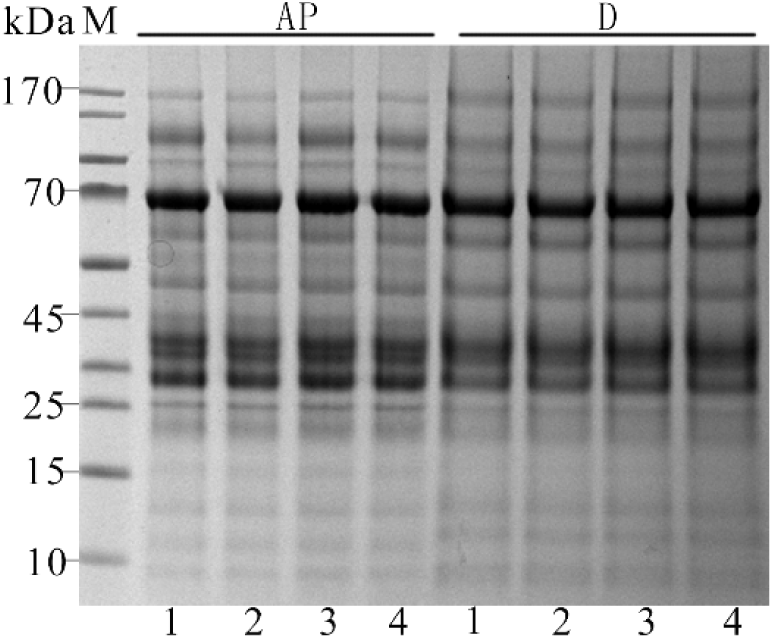
SDS-PAGE analysis of urinary proteins prepared by acetone precipitation and the diaper method. M: marker; AP: urinary proteins prepared by acetone precipitation; D: urinary proteins prepared by the diaper method.

### Evaluating the reproducibility of collection and preservation of urinary proteins using a diaper with label-free quantification

After label-free quantification, 1116 proteins were identified (972, 988, and 969 in each replicate) following three replicates of acetone precipitation. There were 836 shared proteins among the replicates, revealing an 86.1% degree of overlap (Fig. 2A). Simultaneously, 1150 proteins were identified (969, 941, and 964 in each replicate) with 825 shared proteins and an 85.6% degree of overlap (Fig. 2B). In total, 703 proteins were identified following both protocols, revealing an 84.6% degree of overlap (Fig. 2C). Because some differences in protein identification exist between these two methods, future studies should follow only one of the two collection and preservation methods for consistency.

**Figure 2.**
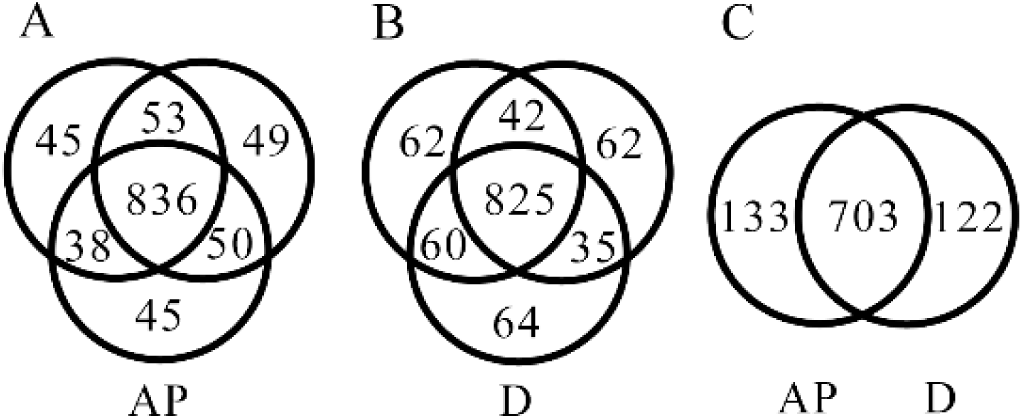
LC-MS/MS identification of urinary proteins prepared by acetone precipitation (AP) and the diaper method (D). A, B: Venn diagrams comparing proteins identified among three independent analyses of the samples purified by acetone precipitation (A) and by diaper method (B). C: Venn diagram comparing proteins identified between these two methods.

To further assess technical reproducibility of the diaper method, we assessed the variation of each identified protein’s abundance among replicates of the two methods (Fig. 4). The average coefficient of variation (CV) was 8.6% for the proteins identified between three replicates following the diaper procedure, and 7.9% for the proteins identified following the acetone precipitation method. For replicates prepared with the diaper procedure, 92.5% of proteins had a CV value < 20%, which is similar to 92.8% for proteins purified with acetone precipitation (Fig. 3). These data suggested that the diaper purification procedure is reproducible.

**Figure 3.**
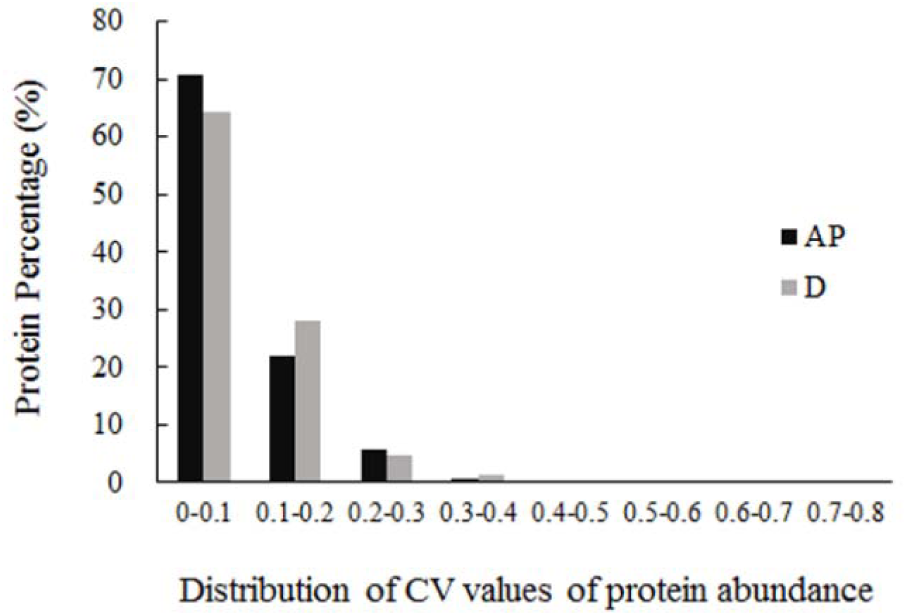
Distribution of CV values of protein abundance by acetone precipitation method (AP) and diaper method (D)

Use of diapers made of fluff pulp facilitates the collection and preservation of urinary proteins from infants, toddlers, and patients with special needs. We also tried using diapers with high percentage superabsorbent polymers; however, eluting proteins from this material was more challenging. Fluff pulp is commonly used, inexpensive, and commercially available, which makes this method economical and suitable for large-scale applications. Theoretically, other materials that absorbs urine but does not absorb proteins may also be used.

## Materials & Methods

### Materials

The absorber (fluff pulp) was purchased from Kotex China (soft cotton, 240 mm). Trypsin was purchased from Promega Company. Iodoacetamide (IAA) and dithiothreitol (DTT) were purchased from Sigma Company. Oasis HLB cartridge was purchased from Waters Company. Nitrocellulose membrane (0.22 μm) was purchased from Millipore Company. Other reagents were of analytical reagent grade.

### Urine collection and preservation

Pooled urine was collected from four volunteers (two males and two females). Written informed consent was obtained from each subject. A workflow explaining the preservation procedure is shown in Fig. 4. The procedure was carried out as follows. (i) The urine sample (30 ml) was poured on a diaper composed of the absorptive fluff pulp and was left to stand for 30 min at room temperature. (ii) Diapers filled with urine were cut, and the fluff pulp was put in a clean beaker, then 80 ml elution buffer (100 mM NH_4_HCO_3_, 50 mM NaCl) was added and mixed with a glass rod. (iii) The mixture was loaded onto the vacuum suction filter bottle described previously (Jia et al., 2014), and the bottle was connected to the vacuum pump causing the solution to pass through the inactivated PVDF membrane (prevents direct contact between fluff pulp and the NC membrane), NC membrane (for protein adsorption), and filter paper layers in a drop-wise manner. (iv) After the proteins were adsorbed onto the NC membrane, the protein-bound membrane was dried using an air dryer or was left to dry at room temperature. (v) The dried membrane was sealed in a vacuum bag and stored at room temperature for two weeks.

**Figure 4.**
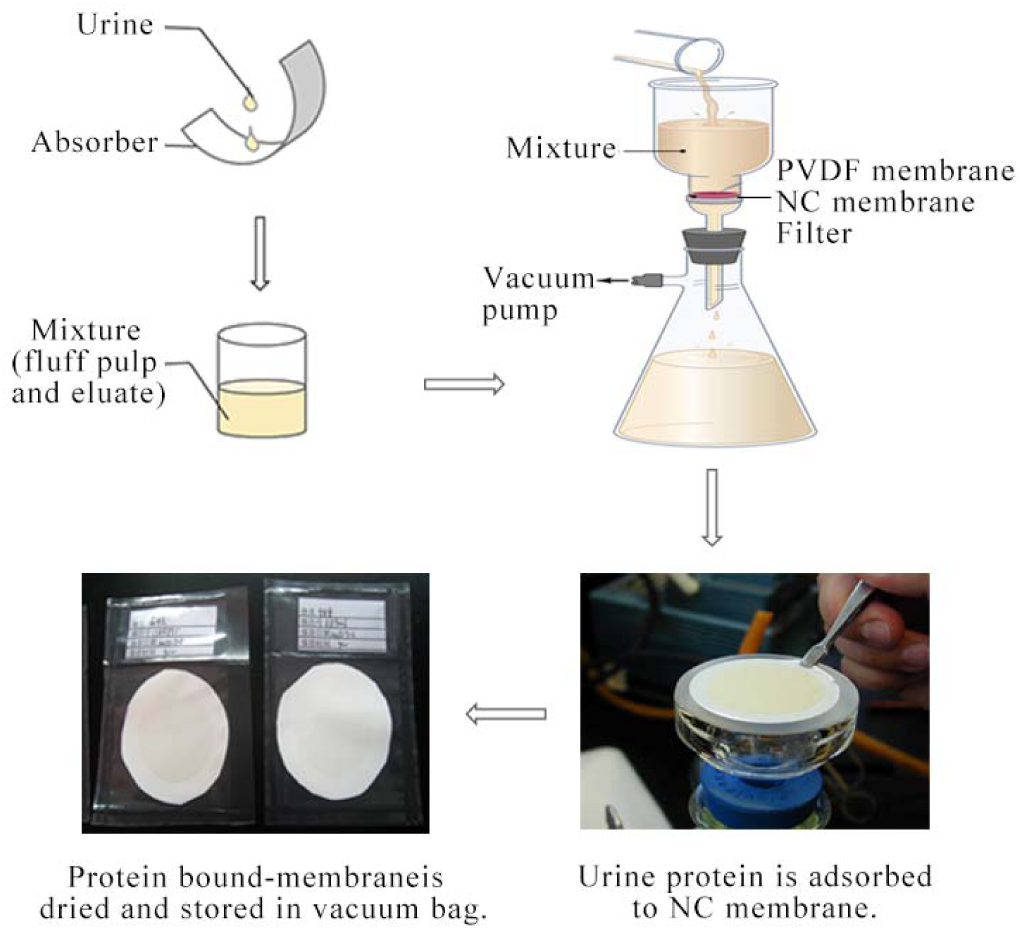
The flow chart of the diaper method (Jia et al., 2014)

### Urinary protein preparation

Urinary proteins were eluted from the membrane using the heating method (Qin and Gao, 2015), and then quantified using the Bradford assay. The pooled urine was centrifuged at 3,500 × *g* for 30 min at 4°C. After discarding the pellet, urinary proteins were extracted from the supernatant by acetone precipitation (Thongboonkerd et al., 2002), and then quantified using the Bradford assay.

### Tryptic digestion

Urinary proteins were digested using filter-aided sample preparation methods (Wisniewski et al., 2009). For digestion, protein solutions were reduced with 4.5 mM DTT for 1 h at 37°C, and then alkylated with 10 mM IAA for 30 min at room temperature in the dark. Finally, proteins were digested with trypsin (1:50) for 14 h at 37°C. The resulting peptides were desalted and then dried using a SpeedVac.

### LC-MS/MS analysis

The digested peptides were dissolved in 0.1% formic acid and loaded on a trap column (75 μm × 2 cm, 3 μm, C18, 100 Å). The eluent was transferred to a reversed-phase analytical column (50 μm × 150 mm, 2 μm, C18, 100 Å) with a Thermo EASY-nLC 1200 HPLC system. Peptides were analyzed with a Fusion Lumos mass spectrometer (Thermo Fisher Scientific). The Fusion Lumos was operated on data-dependent acquisition mode. Survey MS scans were acquired in the Orbitrap using a 350-1550 m/z range with the resolution set to 120,000. The most intense ions per survey scan (top speed mode) were selected for collision-induced dissociation fragmentation, and the resulting fragments were analyzed in the Orbitrap. Dynamic exclusion was employed with a 30 s window. Two technical replicate analyses were performed for each sample.

### Label-free quantitation and statistical analysis

The acquired spectra were loaded to the Progenesis software (version 4.1, Nonlinear, Newcastle upon Tyne, UK) for label-free quantification, as described before (Hauck et al., 2010). Briefly, features with only one charge or more than five charges were excluded from the analyses. For further quantitation, all peptides (with Mascot score > 30 and p < 0.01) of an identified protein were included. Proteins identified by at least one peptide were retained. The MS/MS spectra exported from the Progenesis software were processed with Mascot software (version 2.4.1, Matrix Science, London, UK) in the Swissprot_human database (data 05/03/2013, 20,226 sequences). Search parameters were set as follows: 10 ppm precursor mass tolerance, 0.02 Da fragment mass tolerance, two missed cleavage sites allowed in the trypsin digestion, cysteine carbamidomethylation as fixed modification, and oxidation (M) as variable modifications.

### Compliance and ethics

The author(s) declare that they have no conflict of interest. The consent procedure and this study protocol were approved by the Institutional Review Board (IRB) of the Institute of Basic Medical Sciences, Chinese Academy of Medical Sciences (Project No. 007–2014). In the future, acquisition of human urine samples may not require the same strict regulations as those required for other bio-specimen (Gao, 2015).

## Acknowledgements

This work was supported by the National Key Research and Development Program of China (grant number 2016YFC1306300), the National Basic Research Program of China (grant number 2013CB530850), and funds from Beijing Normal University (11100704, 10300-310421102).

